# AI-readiness Criteria for Biomedical Data in Functional Genomics and Clinical Medicine

**DOI:** 10.1101/2024.10.23.619844

**Authors:** Timothy Clark, Harry Caufield, Jillian A. Parker, Sadnan Al Manir, Edilberto Amorim, James Eddy, Nayoon Gim, Brian Gow, Wesley Goar, Jan N. Hansen, Nomi Harris, Henning Hermjakob, Marcin Joachimiak, Gianna Jordan, Inhee Lee, Maxwell Adam Levinson, Shannon K. McWeeney, Camille Nebeker, Justin Niestroy, Milen Nikolov, Sarah J. Ratcliffe, Justin Reese, Jamie Shaffer, Nathan C. Sheffield, Gloria Sheynkman, James Stevenson, Jake Y. Chen, Chris Mungall, Alex Wagner, Sek Won Kong, Satrajit S. Ghosh, Bhavesh Patel, Andrew E. Williams, Monica C. Munoz-Torres

**Affiliations:** University of Virginia; Lawrence Berkeley National Laboratory; University of California San Diego; University of California San Francisco; Avantiqor; University of Washington; Massachusetts Institute of Technology; Nationwide Children’s Hospital; Stanford University; European Molecular Biology Laboratory - European Bioinformatics Institute; Sage Bionetworks; Oregon Health and Science University; University of Alabama at Birmingham; Boston Children’s Hospital; California Medical Innovations Institute; Tufts University; University of Colorado Anschutz

## Abstract

Biomedical research is rapidly adopting artificial intelligence (AI). The influence of biomedical data preparation on results requires implementing actionable AI-readiness criteria for ethical and explainable AI (XAI) at the "pre-model" stage, encompassing data acquisition, detailed transformations, characterization, and ethical governance. Simple conformance to FAIR (Findable, Accessible, Interoperable, Reusable) Principles is insufficient.

Here, we define criteria and practices for reliable AI-readiness of biomedical data, developed by the **NIH Bridge to Artificial Intelligence (Bridge2AI) Standards Working Group** across seven core dimensions of dataset AI-readiness: **FAIRness, Provenance, Characterization, Pre-model Explainability, Ethics, Sustainability, and Computability.** Conformance to these criteria provides a basis for pre-model scientific rigor and ethical integrity, mitigating downstream risks of bias and error prior to AI modeling. We apply and evaluate these criteria across all four Bridge2AI flagship datasets, spanning functional genomics to clinical medicine, and encode them in machine-actionable metadata bound to the datasets.

This framework, developed against multi-terabyte datasets for reuse in AI applications, sets an empirical benchmark for preparing and evaluating ethical, reusable datasets in biomedical AI.

## Introduction

Artificial intelligence and machine learning (AI/ML) are transforming 21st-century biomedicine, as research now generates massive volumes of high-velocity, multimodal data ^1,2^. Although data are the foundation for AI/ML ^3,4^, adequate standardized definitions of *AI-readiness* for biomedical research remain elusive ^5^. A recent report by the National Academies of Sciences, Engineering, and Medicine (NASEM) identifies this as a critical ‘readiness gap’ in automated research workflows ^6^.

Recent articles highlight ethical data acquisition and pragmatic constraints for AI-readiness ^5,7^. We emphasize here that data are derived by human-made processes - they are constructed ^8^. Therefore transparency in data preparation is a prerequisite for the epistemic validity of data as evidence, as argued by Leonelli ^9^. Without addressing epistemic validity, metadata specifications risk treating complex biological and computational derivations as fictitious ‘ground truths’ devoid of critical context ^8,10^ Data preparation context in metadata is therefore an AI-readiness criterion, whose omission can be harmful in high-stakes biomedical contexts ^11,12^.

Appropriate AI-readiness characterization should establish lifecycle transparency as completely as possible before modeling begins, incorporating metadata on ethical sample and cohort acquisition; instrumentation and post-acquisition processing, including pre-model analytics; sustainable, FAIR data governance ^13^; and computational readiness.

The criteria we present here - seven core dimensions of dataset AI-readiness: (0) FAIRness, (1) Provenance, (2) Characterization, (3) Pre-model Explainability, (4) Ethics, (5) Sustainability, and (6) Computability - are intended to support the current large-scale transition to biomedical AI and should be carefully considered for adaptation and reuse. Dataset descriptions across these criteria must be available as both human- and machine-readable metadata to ensure utility. This framework incorporates lessons learned in the NIH Bridge to AI (Bridge2AI) program, which is designed to produce flagship multi-domain, AI-ready datasets of physiological, molecular, and behavioral data ^14^. Definitions for key terms used in this article follow in Box 1.

### Box 1

**Definitions**

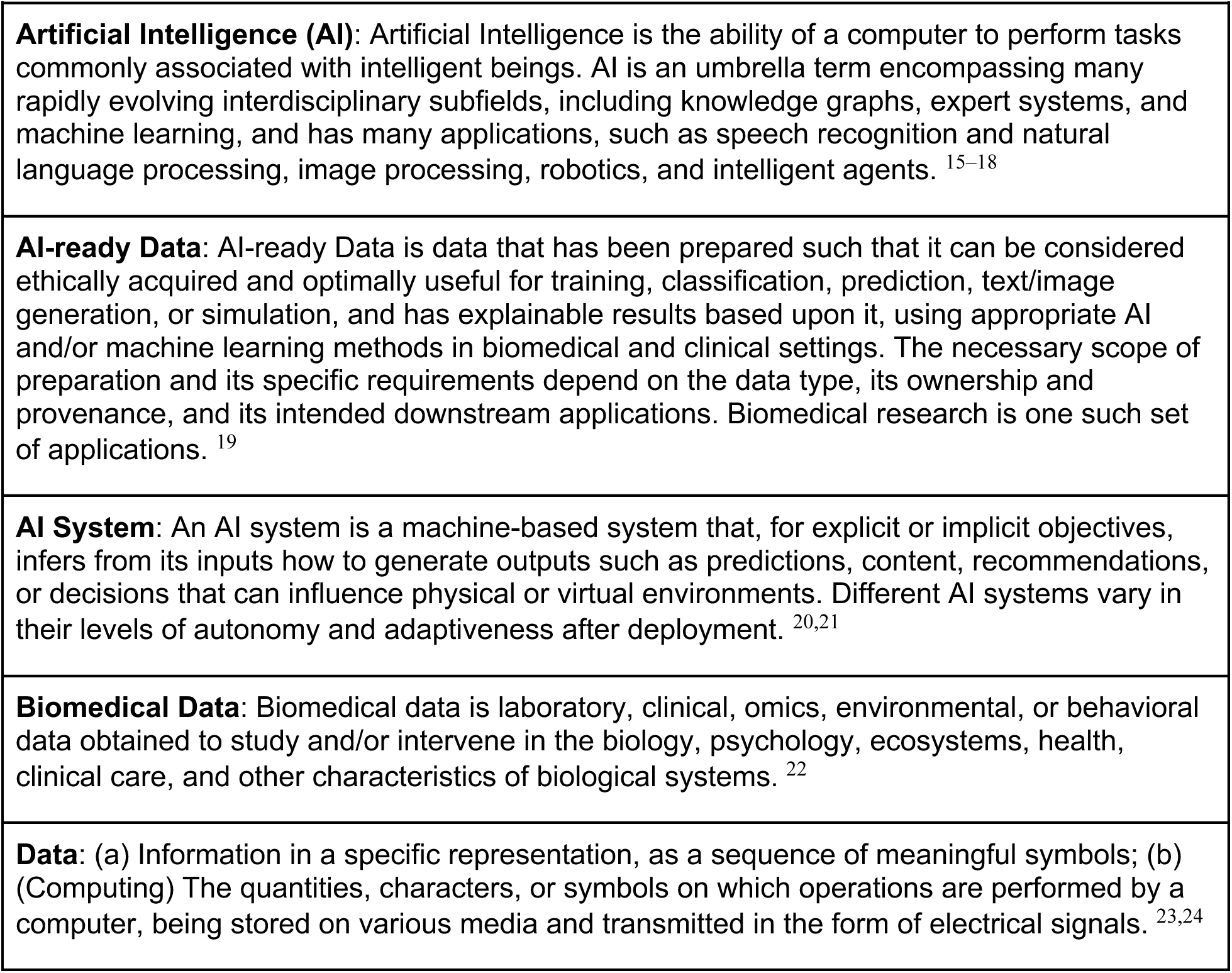
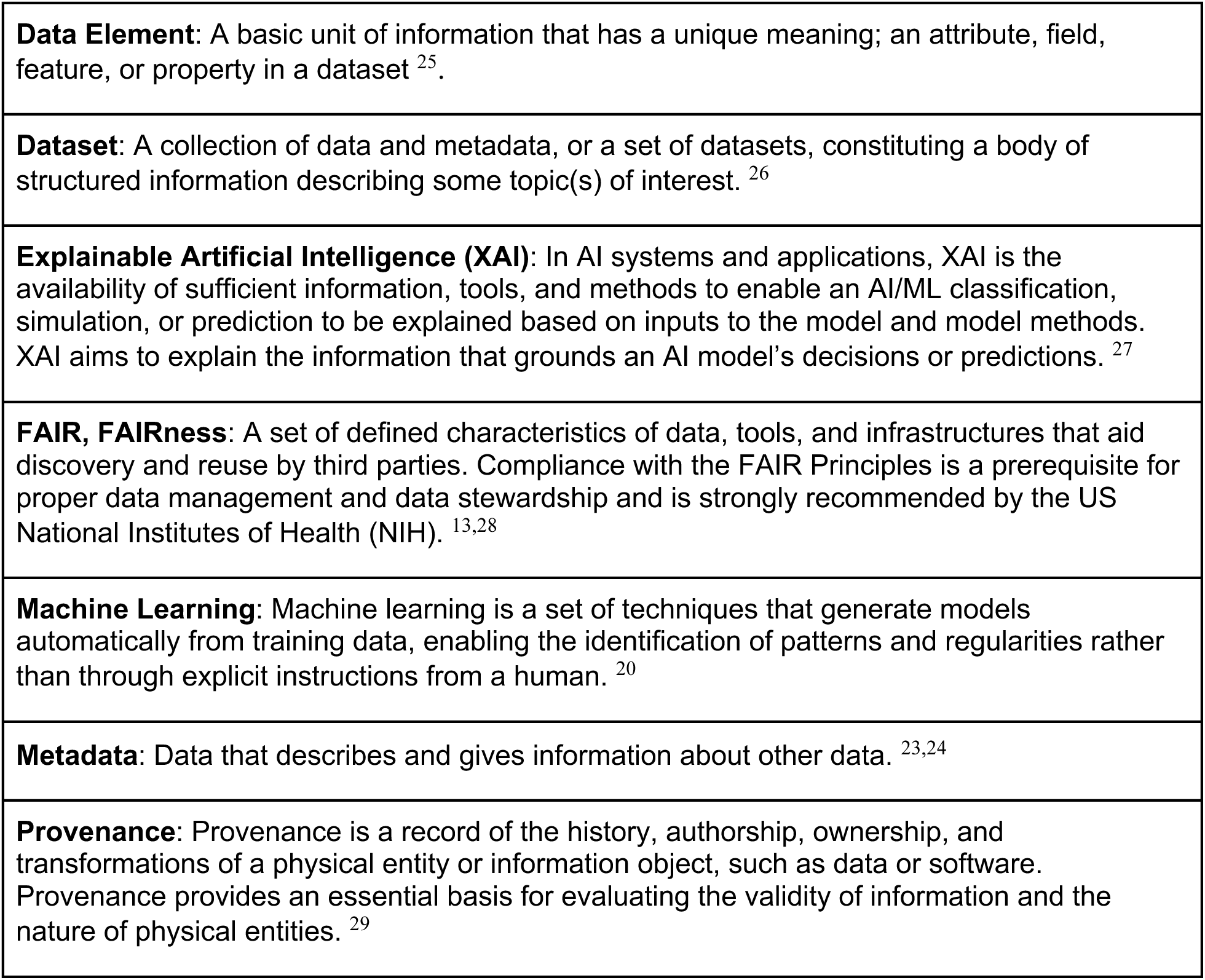

### Approach

The AI-readiness criteria presented here were developed within the Bridge2AI Standards Working Group, comprising experts in AI/ML, genomics, clinical research, data standards, bioethics, ontologies, bioinformatics, and data engineering from across four Grand Challenges (GCs) focused on data production, and the coordinating Bridge Center. These criteria apply to biomedical datasets intended for use in AI/ML training, operations, and evaluation where results will be published in the scientific literature or used for high-impact clinical decisions, and are particularly focused on functional genomics and clinical research use cases.

The Standards Working Group (SWG) coordinated across this large-scale effort comprising more than 40 US and Canadian institutions to prepare these evaluation criteria. The SWG included membership with domain expertise from each of the four GC data generation projects, plus independent standards expertise.

We developed recommendations based on the following inputs:

- Domain Expertise: Ongoing feedback from GC data generation teams preparing data releases.
- Standards: Alignment with established ontologies and NIH metadata guidelines.
- Research Literature: We reviewed extensive literature focused on FAIRness, ethical AI, pre-model explainability, data standards, and data preparedness for AI.
- Validation: Formal ongoing analysis of released Bridge2AI datasets against the proposed criteria.

Our framework builds on the 2019 NIH Advisory Committee to the Director Working Group on AI (ACD AI WG) report ^30^, which identified *Provenance, Description, Accessibility, Sample Size, Multimodality, Perturbations, Longitudinality,* and *Growth as core pillars* of AI readiness. We refined and expanded on these basic ideas to address requirements of preparing large-scale multimodal datasets for AI analysis. Our model stresses, in addition, conformance to ethical and regulatory guidelines for acquisition and governance of data. Biomedical research is guided by regulatory standards including, in the US, HIPAA ^31^ and the Common Rule ^32^; and in the EU, the GDPR ^33^. The Belmont Report ^34^ and the Helsinki Declaration ^35^ are relevant and influential statements on ethics.

Related, foundational work includes the general-purpose MLCommons products Croissant ^36^ and Croissant RAI ^37^, and Health Level Seven (HL7) ^38^. We also adapted concepts from Gebru’s foundational Datasheets for Datasets ^39^ model, from Dataset Nutrition Labels ^40^; and from several other approaches noted later in the text.

A domain-agnostic AI-readiness framework developed by the Open Data Institute (ODI) ^41,42^ in parallel with our 2024 preprint ^43^, provides convergent support for our approach, with 25 of our 28 sub-criteria directly mapping to their dataset, metadata, and infrastructure components ^44^. While the ODI framework article characterizes our model as primarily focused on the surrounding sociotechnical environment, our approach encompasses both the core data infrastructure and this broader context, incorporating distinct gating and software archival criteria. Because data are artifacts of their generating processes, we maintain that integrating this contextual interdependence into foundational metadata is essential for accurately evaluating datasets as scientific evidence ^45–51^.

Our biomedical AI-readiness criteria were empirically grounded in the generation and evaluation of Bridge2AI datasets ^52–55^ over a four-year period. These large-scale resources span functional genomics and clinical medicine, incorporating, in one instance, data from 100,000 patients ^53^, and encompass a diverse array of modalities, including proteomics, genomics, medical imaging, ICU waveforms, wearable sensors, voice recordings, psychometrics, geospatial data, and EHR data. While constructed as foundational, general-purpose datasets to support a wide range of downstream AI applications, they do not negate the need for tailored refinement. We advise data generators to complement our framework by addressing domain-specific clinical nuances and executing task-specific, pre-model data engineering when preparing their datasets for specialized implementations.

### International Standards Context

The International Organization for Standardization and the International Electrotechnical Commission (ISO/IEC) have issued joint AI-related standards dealing with data quality management (DQM), specifying what abstract considerations apply ^56^. Our criteria ask data providers to document precisely what DQM methods were applied, as these are complex and specific to the requirements of functional genomics and clinical data management. Similarly, ISO/IEC 42001 ^57^ recommends that provenance documentation be maintained, but does not specify how to achieve it - which we do specify, using established standards in the W3C PROV family ^58–62^. These include the widely used W3C PROV-O ^58^, and the ISO 23494 Common Provenance Model (CPM) ^61^, a recent ISO standard that extends the W3C PROV-DM provenance data model ^62^ to multi-organizational settings, defining inter-organization connectors where each organization documents only its own provenance segment. It addresses an important use case for interorganizational transfer and we recommend it along with the entire foundational W3C PROV standards family on which it, and our Evidence Graph Ontology (EVI) ^59,60^ are based. CPM packages provenance between organizations and extends PROV-DM; EVI types components within a dataset, and extends PROV-O.

Other ISO/IEC standards on AI govern macro-scale software engineering, organizational, system implementation, and system management, which may be very helpful to organizational AI providers ^63–65^. These issues are important but outside the scope of this article.

Although a comprehensive review of global AI legislation is beyond our scope, the EU AI Act ^21^ is highly relevant as it takes effect for high-risk systems in late 2027 (Annex III high-risk systems) through August 2028 (Annex I product-embedded systems) ^66^. Article 10 mandates the formal documentation and governance of pre-model data preparation, encompassing data annotation, cleaning, and enrichment, and requires frameworks that ensure traceability, dataset integrity, and bias mitigation. While specific compliance standards like prEN 18284 remain in development, emerging EU and ISO standards recommend rigorous data quality management systems without prescribing exact methodologies. Our AI-readiness criteria operate upstream of these forthcoming assessments, providing the foundational governance necessary to satisfy these global regulatory goals.

### Fundamental Requirements of AI-readiness

#### FAIRness, deep provenance, and full characterization

FAIRness is essential for biomedical data; however, the FAIR Principles do not sufficiently account for the intricate derivation histories inherent in biomedical AI applications. We set FAIRness as a Level 0 baseline, but emphasize *Deep Provenance* as a separate requirement.

High-stakes biomedical models, particularly those driving clinical decisions, require full transparency into their processing, beyond simple entity-to-entity assertions. We specify provenance using the W3C Provenance Ontology (PROV-O) and a biomedical-specific extension, the Evidence Graph Ontology (EVI) ^59,60^. EVI provides for deep provenance by decomposition of PROV-O classes into computation-specific terms such as Dataset, Software, Computation, and Annotation. The conventions of science communication were developed to let readers stand as "virtual witnesses" to evaluate how results were produced ^67^. Data in science are now considered as first-class objects ^68^. PROV-O extended by EVI provides a detailed evaluable "methods section" for data resulting from computations, in machine-readable form. Verification taken further, to reproducibility, requires detailed provenance ^69^

Deep provenance ensures that data presented to an AI model derive from verifiable, transparent processes, which must include references to software and, if possible, instrumentation.

Other provenance approaches, such as the Provenance, Authoring, and Versioning (PAV) specification ^70^ or the schema.org *isBasedOn* predicate ^26^, rely on specifying only a particular primary source for a dataset. They do not adequately specify a chain of computations and instrumentation involved in state-to-state transitions.

Widespread irreproducibility of results was identified as a serious issue in 2015 by Begley and Ioannidis who claimed it resulted from nonadherence to good scientific practice ^71^. End-to-end traceability in AI is an important and specific remedy for these issues. It requires that data be resolvable back to their unmodified sources (e.g., raw EHR, NGS reads, imaging, proteomics, sensors, psychometric testing, or clinical trials) through all intermediate steps, including software, models, and computations. If any statistical or computational (e.g., AI/ML) models are used in the processing pipeline, whether for imputation, or for complex inference (as in Bridge2AI Cell Maps for Artificial Intelligence) ^72–74^, they should be described using standard representations equivalent in rigor to Model Cards ^75,76^.

Data must be fully characterized with descriptive metadata on structure, statistical distributions, standards conformance, ethical derivation, biological assumptions, use cases, and inherent biases that may influence model weights. Full provenance graphs in biomedical AI may be constrained to some extent by operational considerations (e.g., *black-box* outsourced laboratory protocols), but we recommend them as goals.

#### Ethics, Regulatory Compliance, and Governance

##### Associated Criteria

These core factors in biomedical AI require verifiable criteria to ensure that data are well-governed and ethically and legally compliant. We define criteria in this section to ensure that (1) the intended applications align with consent and institutional agreements; (2) governance attributes including Data Use Agreements (DUA) are represented in machine-readable format; (3) socio-technical context of data collection, whether directly from individuals in a cohort, or from banked samples, is represented to make potential sources of bias transparent.

##### Human-mediated Oversight

Our criteria provide for human-mediated oversight. We require documentation of individuals responsible for ethics review, IRB and protocol-specific authorization, and designation of a data governance chairperson. Depending upon project scale, governance may be overseen by a Principal Investigator or by a well-organized Governance Committee. The responsible governance chair and ethics contact will ensure appropriate governance of the project, supervise alignment with Belmont, Menlo, and CARE principles of ethical guidance ^34,77–79^, and other standards relevant to protection of subjects and privacy. The governance chair is the point of contact and certifies all data releases.

##### Ethical Licensing

We require that explicit license terms be identified in metadata, as appropriate to regulatory requirements, using a granular Data Use Agreement or common licenses, such as Creative Commons CC-BY and its variants ^80^.

#### Sustainability

Sustainability ensures that the compute-intensive investment in AI training remains reproducible through long-term archiving in trusted, sustainable repositories ^81^. For controlled-access sensitive data repositories, additional security measures will apply to ensure sound security practices, in accordance with applicable government regulations and funding agency requirements. We require that the chosen repository supports long-term persistence of data and metadata packages with project-appropriate security levels.

#### Pre-model Explainability

Pre-model Explainability is the transparency and reliability summit of our criteria, summarizing metadata supporting all other criteria into a human- and machine-readable Datasheet and providing cryptographic hash integrity seals for the metadata plus data packaging. Our Datasheets substantially extend the well-known Gebru et al. (2021) Datasheets for Datasets model ^39^, making them both actionable and accessible through an architecture for the specification and analysis of these metadata ^82,83^. Developing this framework was a central challenge of our project, requiring a consensus-driven approach among domain scientists, computational biologists, clinicians, and ontologists, each of whom was previously committed to distinct modeling frameworks. Our framework uses a middle exchange layer to transform heterogeneous, domain-specific data into standard interoperable packages.

This architecture includes LinkML metadata ^84^ along with the provenance graphs in a community-standard, lightweight RO-Crate-based Exchange Layer ^82^. LinkML provides a basis for later deep semantic enrichment. We also provide software for the evaluation of criterion conformance and its visual representation in human-readable datasheets ^85,868081828385^. We maintain close collaboration with the RO-Crate center of practice at the University of Manchester and with the LinkML community (many of whom are in the Bridge2AI team) to ensure continued harmonization and integration of these approaches.

Although we provide supporting tools ^87^ to facilitate the implementation of this metadata model, the framework itself is platform-agnostic and does not strictly require their use.

The overall purpose of our criteria remains the same: it is to enable managed self-improvement of AI-readiness for datasets intended for publication in the scientific record, or to be used in clinical evaluations. The criteria establish targets for maximum pre-model transparency.

### Computability

Basic requirements for computability are use of standard data formats, data accessibility, computing resource requirements, split or holdout description, and exemplars of input data. Standardized well-understood data formats are needed so that binary information can be mapped to intended meaning. Computational accessibility via standard exchange protocols or documented APIs is required simply to access the data. If any specific resources are required, metadata must document them. Split or holdout characterizations of training, test, and validation datasets help prevent overfitting and data leakage, while providing guidance for hyperparameter tuning. Examples of data inputs help in understanding outliers and ensure they are not hidden by selection statistics.

### Dimensions of AI-readiness

AI-readiness is a dynamic, context-dependent developmental property of specific data sets. We score it along multiple dimensions based on readiness scores for major components, yielding a characteristic multidimensional score matrix. Achieving AI-readiness in any particular use case is a collaborative, developmental, research-driven task ^19,88,89^.

AI-readiness, as defined here, extends beyond utility, convenience, or tractability for computer scientists and informaticians. It emphasizes the reuse of data and the generation of results that are ethical, scientifically valid, explainable, interpretable, and sustainable. Our criteria for AI-readiness are a framework for assessing progress toward these goals. We consider scientific validity and research integrity as ethical practices ^90,91^. Ultimately, our principal goal is data that are available, reusable, deeply characterized, standardized where possible, and that provide foundational support for ethical explainability of results. These criteria are meant as a foundation for reuse and adaptation. By adapting these criteria, research communities can iteratively build toward fully responsible AI practices.

### AI-readiness Criteria

#### Fundamental Criteria

AI-readiness imposes properties that extend far beyond basic FAIR compliance, requiring more rigorous specification and novel metadata extensions.

AI-readiness implies that data must be: **FAIR, Provenanced, Characterized, Ethical, Pre-model Explainable, Sustainable, and Computable**. As illustrated in the AI-readiness Wheel (**Fig. 1**), these seven dimensions are not isolated silos; rather, they form an interdependent ecosystem in which the strength of one, such as Deep Characterization, directly enables the success of another, such as Pre-model Explainability.

**Figure 1.**
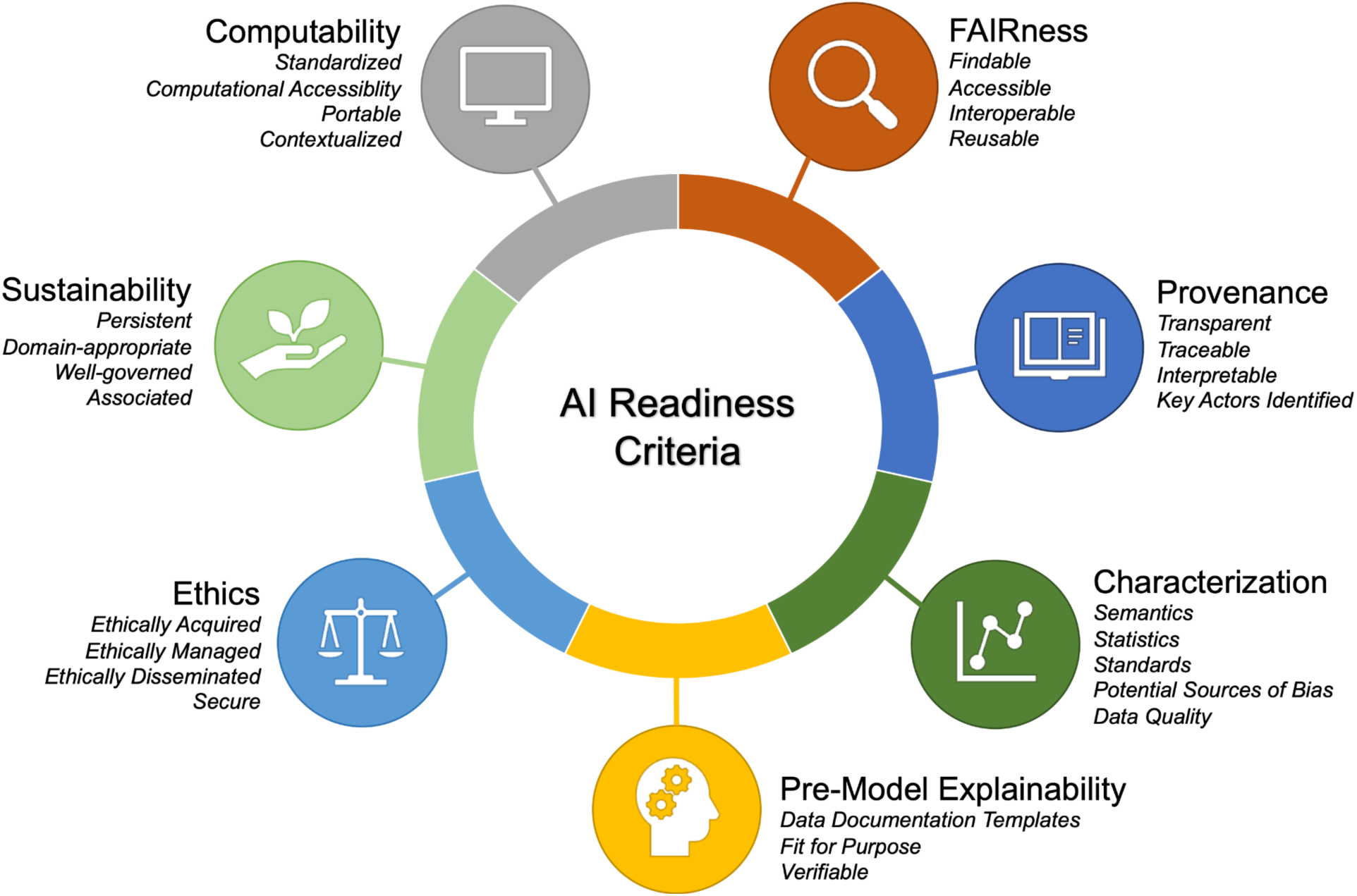
AI-readiness Wheel. Seven major criteria of AI-readiness were developed for Bridge2AI datasets, along with their relevant sub-criteria (*italics*) as detailed in Table 1.

**Table 1.** AI-readiness Criteria, Sub-criteria, and Practices. The asterisk (*) indicates that this criterion or sub-criterion must be satisfactorily fulfilled (“gating”).

| **ID** | **Criterion** | **Practice<sup>†</sup>** | **Suggested resources** |
|---|---|---|---|
| 0 | FAIRness* |  |  |
| 0.a | Findable | Deposit datasets in a searchable FAIR-compliant data repository. | FAIRsharing.org<sup>92</sup><br>NIH GREI repositories<sup>93</sup> |
| 0.b | Accessible | Descriptive metadata should always be available and accessible, even if the dataset is restricted, unavailable, or de-accessioned. Ensure metadata conforms to standards such as DCAT (Data Catalog Vocabulary), DataCite schema, and/or schema.org. | DCAT<sup>94</sup><br>schema.org<sup>95</sup><br>DataCite schema<sup>96</sup> |
| 0.c | Interoperable | Wherever possible, provide data and metadata using formally defined specifications for digital objects. | RDF<sup>97</sup><br>JSON-LD<sup>98</sup> |
| 0.d | Reusable | Attach a clear and accessible data usage license that allows the responsible use of AI/ML applications or link to a Data Use Agreement (DUA). | Creative Commons<sup>80</sup> |
| 1 | Provenance |  |  |
| 1.a | Transparent* | Identify primary data sources, e.g., clinical data from EHR at a given hospital, clinical trials data, or laboratory data from an instrument. | W3C PROV<sup>29</sup>, EVI<sup>59,60</sup>, CPM<sup>61</sup>, PAV<sup>70</sup>, FAIA<sup>99</sup><br>OMOP<sup>100</sup>, RRID<sup>101</sup> |
| 1.b | Traceable | Identify key data transformation steps, with links to software, at an appropriate level of detail, using a machine-readable representation. | W3C PROV<sup>29</sup>, EVI<sup>59,60</sup>, CPM<sup>61</sup> |
| 1.c | Interpretable | Make software for key data transformation and analysis steps available in a sustainable repository for transition steps interpretability. | Zenodo<sup>102–104</sup><br>Software Heritage<sup>105</sup><br>GitHub<sup>106</sup> |
| 1.d | Key Actors Identified | Identify people and organizations responsible for obtaining and processing the data, the samples, and the subject groups from which the data were produced. Reference these parties along with other dataset metadata. | ORCID<sup>107</sup><br>ROR<sup>108</sup> |
| 2 | Characterization |  |  |
| 2.a | Semantics | Use comprehensive descriptive metadata for datasets, including a detailed abstract, dataset keywords, and subject-specific vocabularies (e.g., MeSH) to enable detailed search and discovery. | DataCite schema<sup>96</sup> |
| 2.b | Statistics | Provide domain-appropriate statistical characterizations of key features of the dataset to inform analysis planning. Ensure missing values are encoded consistently. | Pandas/Polars<sup>109,110</sup><br>Frictionless Python Framework<sup>111</sup> |
| 2.c | Standards* | Provide a machine-readable data dictionary or schema for each dataset, linked to the dataset metadata, and referencing any relevant standards. | Frictionless Data<sup>111</sup><br>JSON Schema<sup>112</sup><br>Pydantic<sup>113</sup> |
| 2.d | Potential Sources of Bias | Describe known sources of bias in the data and assumptions made in collecting, processing, or interpreting the data. Include any known explanations regarding missing values, including reasons for missingness, and the degree to which data represents a state vs. a control (e.g., disease state vs. healthy state). | Croissant RAI<sup>37</sup> |
| 2.e | Data Quality | Describe and/or link to data quality control procedures. | W3C DQV<sup>114</sup> |
| 3 | Pre-model Explainability |  |  |
| 3.a | Data Documentation Template | Machine-readable metadata and a linked human-readable document should support a domain-appropriate extension of the original Gebru Datasheets concept for pre-model explainability, as detailed in the present article. Healthsheets information may be a helpful supplement. | Datasheets<sup>39</sup><br>Health Sheets<sup>115</sup> |
| 3.b | Fit for Purpose | Identify appropriate and inappropriate use cases for the data set. Link to any previously published analyses using this data. | Croissant RAI<sup>37</sup><br>DUO<sup>116</sup> |
| 3.c | Verifiable | Provide a mechanism for ensuring the integrity of each raw or processed dataset, such as a cryptographic hash. | SHA256<sup>117</sup><br>Merkle Trees<sup>118</sup> |
| 4 | Ethics |  |  |
| 4.a | Ethically Acquired* | Describe ethical data acquisition consistent with accepted principles (e.g., Belmont Report, Menlo Report, CARE Principles) sufficient to evaluate for intended use, along with a management plan. | Belmont Report<sup>34,77</sup><br>Menlo Report<sup>78</sup><br>CARE Principles<sup>79</sup> |
| 4.b | Ethically Managed | Data management must align with ethical principles throughout the health AI lifecycle. Indicate privacy-protection processing, if any, sufficient to evaluate ethical status for intended use. | HL7 privacy protection metadata<sup>119</sup> |
| 4.c | Ethically Disseminated | Specify a licensing agreement and/or data use agreement (DUA) on as open terms as ethical and sustainability considerations permit. Specify contact information for a data access committee chair and for an ethics reviewer. | Creative Commons<sup>80</sup><br>GA4GH DACRes<sup>120</sup> |
| 4.d | Secure | Specify security requirements for storing and accessing this data, e.g., “public,” “controlled access only,” etc. | HL7 privacy protection<sup>121</sup> |
| 5 | Sustainability |  |  |
| 5.a | Persistent | Ensure unprocessed data is preserved in an archive that complies with privacy laws and retention guidelines, enabling future reprocessing and publishing of revised data. | TRUST Principles<sup>81</sup> |
| 5.b | Domain-appropriate | Ensure single-domain raw or processed data are deposited in a FAIR-domain-appropriate specialist repository if available. | NIH GREI repositories<sup>93</sup><br>FAIRsharing.org<sup>92</sup> |
| 5.c | Well-governed | Implement a data governance model before data capture, including a repository that facilitates data stewardship in the future and governance that accounts for maintenance, terms, policy changes, and fairness. | FAIRsharing.org<sup>92</sup><br>NIH GREI repositories<sup>93</sup> |
| 5.d | Associated | Document project-level connections between data components and elements in a machine-readable manner. | Croissant<sup>36</sup><br>PROV-O<sup>58</sup><br>Frictionless Data<sup>111</sup><br>RO-Crate<sup>122</sup> |
| 6 | Computability |  |  |
| 6.a | Standardized | Datasets must adhere to established, documented standards, including metadata standards, and their adherence can be validated deterministically. | Croissant<sup>36</sup><br>Croissant RAI<sup>37</sup> |
| 6.b | Computationally Accessible | Provide a mechanism to access data through established exchange protocols or a well-documented API. | FAIRsharing.org<sup>92</sup><br>NIH GREI-participating repositories<sup>93</sup> |
| 6.c | Portable | Maximize portability across computational resources where possible. If data use requires specific resources, provide machine-readable documentation defining these resources. | CodeMeta<sup>123</sup> |
| 6.d | Contextualized | Include any considerations regarding splits of the data, including any information withheld at any point of data collection and processing. Provide examples of data components to aid understanding of their general structure and content. | Croissant<sup>36</sup> |

<sup>†</sup> Practices may impact multiple criteria; the most relevant relationship is shown for brevity

#### Detailed AI-readiness Criteria

To provide more precise implementation guidance, we developed the following detailed criteria and supporting practices. We have reviewed these practices against a related effort in another domain, Earth and Space Sciences, as a consistency check ^89^. **Table 1** describes AI-readiness criteria and relevant practices that support their achievement. These criteria are the first set of multidisciplinary consensus AI-readiness criteria derived across multiple biomedical projects of this scale: forty institutions, multi-terabyte data from 100,000 patients, large-scale functional genomics and proteomics, wearable sensor, geolocation, and environmental data, continuous patient monitoring, imaging, voice spectrograms, and Electronic Health Records (EHR) data.

The AI-readiness of biomedical data can be evaluated against machine-readable metadata using these criteria. We provide evaluation radar plots against these criteria on interim datasets of the four large-scale Grand Challenge (GC) programs in Bridge2AI in **Figure 2**, along with tools to assist scoring based on metadata in the RO-Crate dataset packages ^52–55^. The tools provided include an evaluation rubric ^124^ with detailed scoring assistance ^85^, to evaluate each sub-criterion. These are scored on a three-point ordinal rating scale and we then calculate a proportional score for each major axis. Scoring detail for each GC is shown in Table 2.

**Figure 2.**
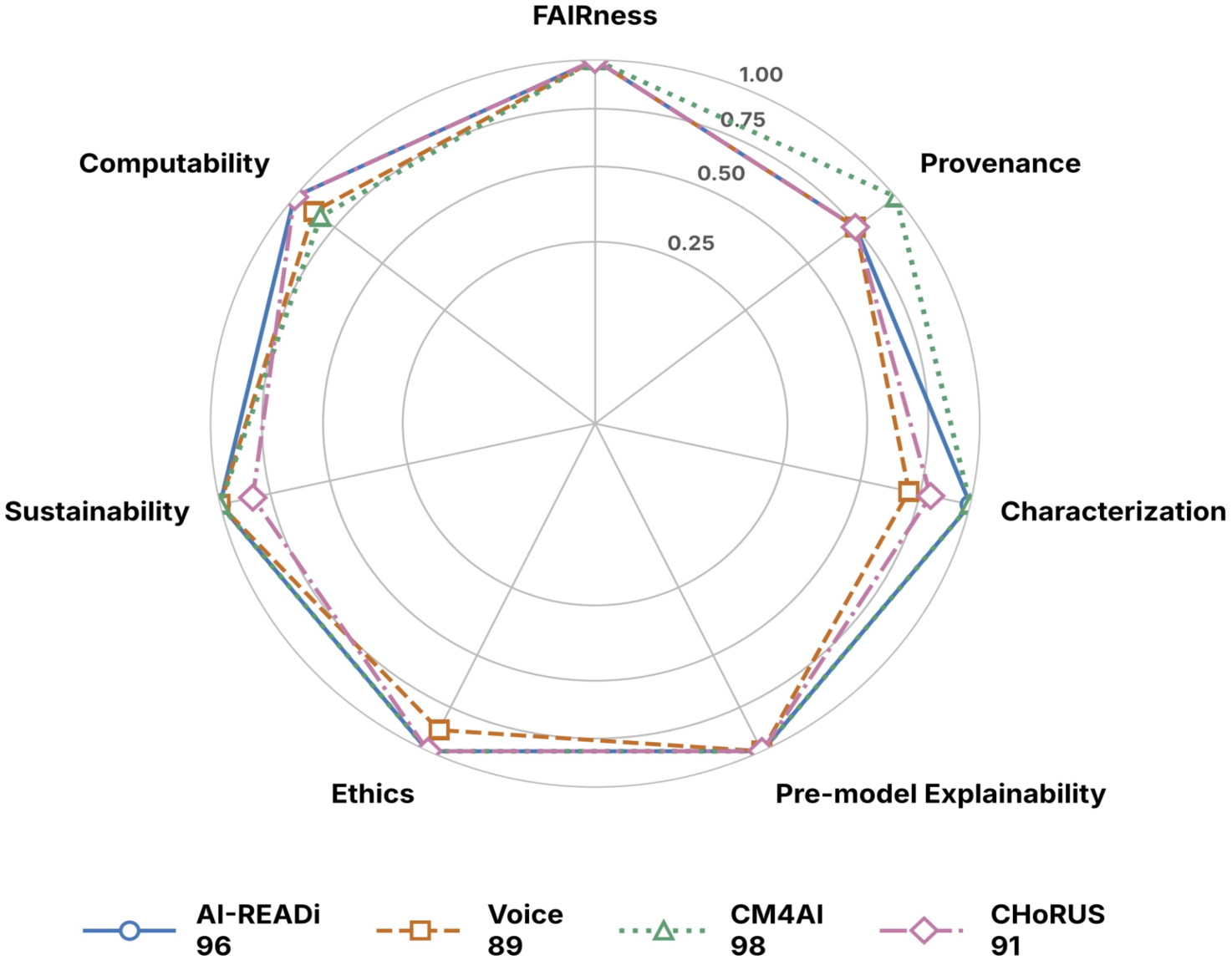
AI-readiness of near-final releases of the four Bridge2AI Grand Challenge datasets. ^52–55^. Scoring is a per-axis proportion of maximum score, omitting any N/A sub-criteria. Radial axes are square root scaled. Total score is 100 times the average of axis scores. If any axis had failed a gate, it would have been marked “Gating FAIL” and that label carried up to the total alongside the score. This evaluative framework provides feedback to guide metadata and data improvement on readiness axes where the scores are weaker, throughout the release lifecycle. It also sets a foundation for adjusting scoring weights for specialized scientific domains or use cases.

**Table 2.**
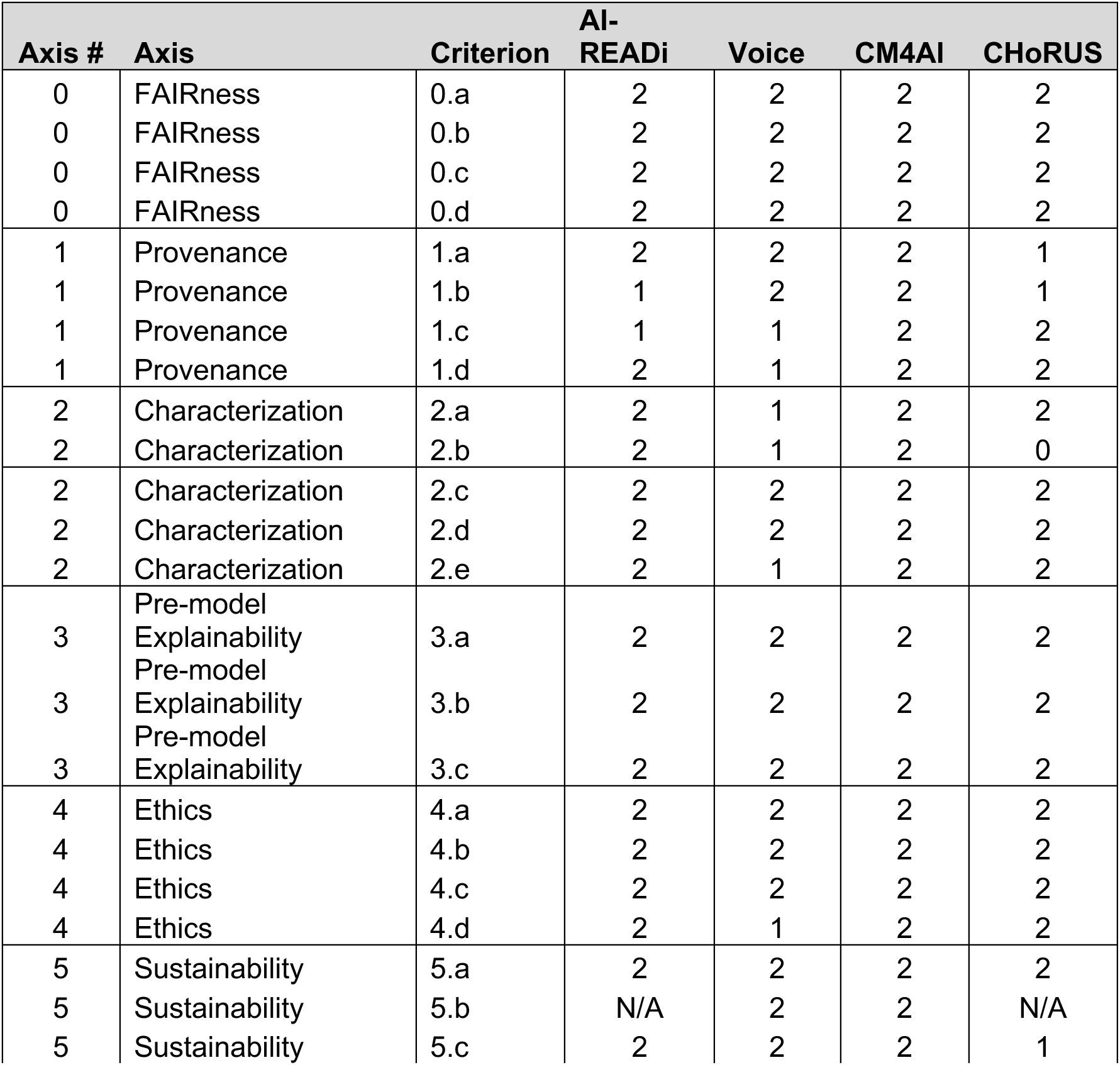
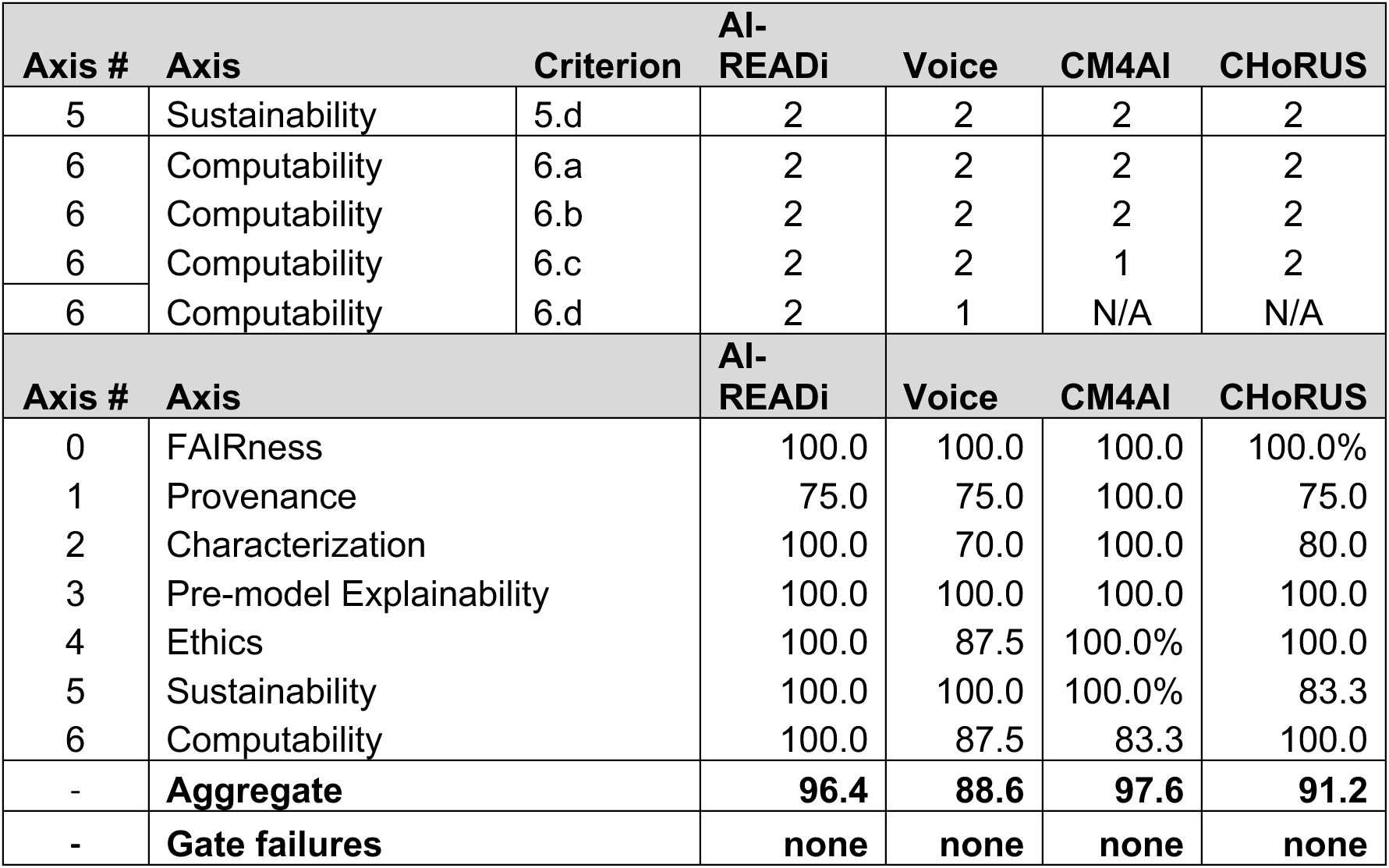
AI-readiness detailed scores. Our rubric provides sub-criterion scores according to answers provided about the data and metadata by representatives from each GC in collaboration with Bridge Center investigators. Bridge Center investigators made the final score assignments. A rating calculation computes axis scores across sub-criteria, using the method described in the Figure 2 caption. All GCs scored above 88. Weaker scores reflect areas for improvement in developing the final data releases.

| Axis # | Axis | Criterion | AI-READi | Voice | CM4AI | CHoRUS |
|---|---|---|---|---|---|---|
| 0 | FAIRness | 0.a | 2 | 2 | 2 | 2 |
| 0 | FAIRness | 0.b | 2 | 2 | 2 | 2 |
| 0 | FAIRness | 0.c | 2 | 2 | 2 | 2 |
| 0 | FAIRness | 0.d | 2 | 2 | 2 | 2 |
| 1 | Provenance | 1.a | 2 | 2 | 2 | 1 |
| 1 | Provenance | 1.b | 1 | 2 | 2 | 1 |
| 1 | Provenance | 1.c | 1 | 1 | 2 | 2 |
| 1 | Provenance | 1.d | 2 | 1 | 2 | 2 |
| 2 | Characterization | 2.a | 2 | 1 | 2 | 2 |
| 2 | Characterization | 2.b | 2 | 1 | 2 | 0 |
| 2 | Characterization | 2.c | 2 | 2 | 2 | 2 |
| 2 | Characterization | 2.d | 2 | 2 | 2 | 2 |
| 2 | Characterization | 2.e | 2 | 1 | 2 | 2 |
| 3 | Pre-model Explainability | 3.a | 2 | 2 | 2 | 2 |
| 3 | Pre-model Explainability | 3.b | 2 | 2 | 2 | 2 |
| 3 | Pre-model Explainability | 3.c | 2 | 2 | 2 | 2 |
| 4 | Ethics | 4.a | 2 | 2 | 2 | 2 |
| 4 | Ethics | 4.b | 2 | 2 | 2 | 2 |
| 4 | Ethics | 4.c | 2 | 2 | 2 | 2 |
| 4 | Ethics | 4.d | 2 | 1 | 2 | 2 |
| 5 | Sustainability | 5.a | 2 | 2 | 2 | 2 |
| 5 | Sustainability | 5.b | N/A | 2 | 2 | N/A |
| 5 | Sustainability | 5.c | 2 | 2 | 2 | 1 |
| **Axis #** | **Axis** | **Criterion** | **AI-READi** | **Voice** | **CM4AI** | **CHoRUS** |
| 5 | Sustainability | 5.d | 2 | 2 | 2 | 2 |
| 6 | Computability | 6.a | 2 | 2 | 2 | 2 |
| 6 | Computability | 6.b | 2 | 2 | 2 | 2 |
| 6 | Computability | 6.c | 2 | 2 | 1 | 2 |
| 6 | Computability | 6.d | 2 | 1 | N/A | N/A |
| **Axis #** | **Axis** |  | **AI-READi** | **Voice** | **CM4AI** | **CHoRUS** |
| 0 | FAIRness |  | 100.0 | 100.0 | 100.0 | 100.0% |
| 1 | Provenance |  | 75.0 | 75.0 | 100.0 | 75.0 |
| 2 | Characterization |  | 100.0 | 70.0 | 100.0 | 80.0 |
| 3 | Pre-model Explainability |  | 100.0 | 100.0 | 100.0 | 100.0 |
| 4 | Ethics |  | 100.0 | 87.5 | 100.0% | 100.0 |
| 5 | Sustainability |  | 100.0 | 100.0 | 100.0% | 83.3 |
| 6 | Computability |  | 100.0 | 87.5 | 83.3 | 100.0 |
| - | **Aggregate** |  | **96.4** | **88.6** | **97.6** | **91.2** |
| **-** | **Gate failures** |  | **none** | **none** | **none** | **none** |

All scores were human assigned. The criteria and sub-criteria are unweighted. Use-case applicability considerations may dictate weighting these sub-criteria, while keeping in mind that the mandatory "gating" criteria and sub-criteria (FAIRness 0.a-0.d, Provenance 1.a, Characterization 2.c, and Ethics 4.a) establish a pass/fail threshold for the entire dataset. Grounded in machine-readable AI-readiness metadata, this framework allows for the generation of human-readable Datasheets that expand on the concepts of Gebru et al. ^39^ while providing the transparency necessary to support human review, accommodate variations in metadata structures, and enable other users to easily apply scoring. Detailed scores assigned by human reviewers from each GC in consultation with the Bridge Center are presented in Table 2.

We emphasize that assessment of these readiness criteria is a self-improvement exercise, for which self-evaluation will likely be the norm. No current standards body or evaluation center is in place to grade conformance independently, although large-scale programs are encouraged to fund this type of activity. Use of transparent rubrics and scoring criteria for AI-readiness provides an important degree of openness for review by the community, where the datasets are open or accessible under reasonable data access agreements.

#### Translating AI-readiness Criteria into Exchangeable Metadata

The AI-readiness criteria described in this article help support data reuse and evaluation across heterogeneous biomedical and computational contexts. To enable practical reuse, these criteria must be operationalized as structured, exchangeable metadata describing the datasets along the major criteria axes. Within Bridge2AI, we translate the criteria into machine-actionable metadata wrapped in standard, lightweight RO-Crate version 1.2 packages ^125^. This approach employs widely adopted serializations and vocabularies for interoperability across domains. Bridge2AI RO-Crate packages provide metadata using: JSON-LD graphs and the schema.org vocabulary for semantic content ^95,98^; JSON Schema dataset structure descriptions ^112^; and Frictionless Data schema evaluation ^111^. Provenance information is represented using the W3C PROV data model as a structural backbone, supplemented by EVI profiles ^59,60,29^. As established earlier, EVI is a domain-specific extension of PROV-O ^58^ that provides biomedical specializations of the core PROV classes (Entity, Activity, and Agent). Dual use of PROV-O and EVI preserves compatibility with existing provenance-aware tools and infrastructures while enabling more semantically precise biomedical domain descriptions and epistemic justification. Bridge2AI FAIRSCAPE tools ^86^ generate RO-Crate packages conforming to this model and can optionally translate these packages to LinkML semantic modeling language for deeper semantic integration across projects when required.

RO-Crate packages may be generated through many types of workflows and validated at varying levels of rigor. While tooling such as Bridge2AI’s FAIRSCAPE^87^ provides one concrete implementation of this approach, the translation of AI-readiness criteria into exchangeable metadata is not dependent on any specific software system and is intended to be broadly applicable across biomedical data ecosystems.

#### Challenges and Limitations

These criteria are meant for self-improvement of AI-readiness, not as a conformance standard. The Bridge2AI datasets were evaluated collaboratively by personnel from the Bridge Center Standards Working Group, the CM4AI tools developers, and the GCs. Readers may assess the scores for themselves, using the data we provide here, as well as the adequacy of our rubrics and scoring methods. It remains to be established how far these criteria may be generalized through adoption and adaptation outside Bridge2AI. Even where a large program has funding for “independent” evaluators, conflicts of interest may occur, so we recommend relying on transparency of scoring methods, data, and metadata. Highly descriptive metadata along all seven criteria is the basic requirement for this assessment. Scores measure metadata as supplied.

Preparation of AI-ready biomedical datasets imposes challenges, requiring additional effort beyond simply capturing measurements or observations for statistical analysis. This becomes more complex when effort increases as datasets are meant to meet multiple use cases and to be sustainable over time, as in Bridge2AI, rather than addressing one-off, highly focused research questions. For use-case-specific studies, additional data engineering steps, such as feature engineering, may need to be added for context specificity. Those added steps will need to be documented in metadata – including the provenance graph – at the same level as the rest of the dataset metadata.

AI-readiness data preparation requires a deep understanding of the data, the predictive task(s) for which the data will or may be used, the scientific domain of the data, statistical methods, AI technologies, biomedical data standards, and appropriate ethical practices. It also requires at least some attention, depending upon the project scope and intended longevity, to sustainability within the biomedical data ecosystem. There is a tension between recording practices in machine-readable metadata, and the agility of typical computational experimentation. It is the task of effective program and project leadership to navigate this tension successfully without compromise to metadata clarity and interpretability. Our approach particularly respects the need for pre-model explainability by clearly defining provenance using four unique, stand-alone sub-criteria (1.a-1.d; see **Table 1**).

In clinical studies, the ethical treatment of human subjects and the protection of their data are major concerns. It requires close attention to proper privacy-preserving practices, and responsible data stewardship, further emphasizing the need for the Provenance and Ethics criteria to ensure that data-use limitations, compliance, intellectual property, and other restrictions are clearly stated and adhered to in downstream data use.

Certain inherent limitations are implied by the time-, place-, technology-, and culture-boundedness of our efforts. AI/ML applications and capabilities are a rapidly progressing, revolutionary scientific and societal development. Our understanding of data ethics and society’s ability to democratically control and adapt AI technologies for the widest possible social benefit must surely evolve. Cultural, ethnic, and gender definitions used in these datasets today may seem archaic in ten or twenty years, and what we do not conceive of as biases today may seem biased tomorrow. Thus, best practices must continue to evolve alongside the field of biomedical AI/ML.

A final limitation is the reliance on the US- and EU-focused legal and standards environment. We did not assess AI-readiness legal requirements outside these contexts, and moving beyond them will impose new challenges. However, our work is consistent with and supports the basic requirements for transparency, accountability, and inclusiveness described in *Ethics and governance of artificial intelligence for health: WHO guidance* ^126^, which are transnational in scope.

### Conclusions and Future Directions

AI-ready biomedical data preparation and evaluation require practices to establish data **FAIRness**, **Provenance**, **and Characterization**, with support for **Pre-model Explainability**, **Ethics, Sustainability**, and **Computability**. These should be reflected in the human- and machine-readable metadata associated with an AI-ready dataset. We have outlined criteria reflecting our recommended practices and multiple methods to evaluate adherence to these criteria. The criteria proposed here are currently used in the Bridge2AI program and have evolved across multiple data releases through intensive discussions among multi-domain experts and in parallel with metadata packaging tools. Developing readiness criteria in parallel with major data resources enabled our participating investigators to deeply examine requirements for AI-readiness. It prompted passionate methodological discussions among experts and ultimately motivated the criteria and evaluation methods described in this article. These criteria, adapted to varying circumstances, with adherence to the gating criteria and appropriate weighting of the non-gating criteria according to data modality, intended use, regulatory setting, and potential patient risk, can enable biomedical AI that produces ethical, reliable, transparent, and explainable results. Our major contributions outlined in this article include:

- Defined practices and criteria for AI-readiness of biomedical data;
- A formal evaluation approach against these criteria;
- Detailed evaluation of the Bridge2AI Grand Challenge datasets.

Some open-source tools supporting AI-readiness developed in Bridge2AI include:

- RO-Crate structures for multimodal data with Deep Provenance;
- Extended Evidence Graph Ontology (EVI) extension of W3C PROV;
- JSON Schema, Pydantic, and LinkML models for AI-readiness Datasheets;
- Rubric and scoring model for dataset evaluation against AI-readiness evaluation criteria; and
- The FAIRSCAPE AI-readiness framework.

These tools, though not required to implement the AI-readiness criteria, will continue to be developed and fully extended across all Bridge2AI program components.

We invite feedback on this work and collaboration with the broader biomedical AI/ML community, including users of Bridge2AI datasets. The perspectives presented here reflect the experience of teams producing datasets intended to serve as flagship examples of best practices. As these data are used to develop effective AI methods, their application will help identify strengths of the proposed approach and guide future improvement. We encourage the use of these metrics to improve AI-readiness across data products and to assess the suitability of reused datasets.

## Data and Software Availability Statement

### Datasets

The four Bridge2AI Grand Challenge datasets cited below were evaluated against the AI-readiness criteria using a detailed rubric and scoring methodology..

`o` Clark T, Parker J, Al Manir S, et al. Cell Maps for Artificial Intelligence - June 2026 Data Release (Beta). Published online July 15, 2026. doi:10.18130/V3/HIGT4C ^52^
`o` Rosenthal E, Kamaleswaran R, Levites Strekalova YA, Williams A, Cordes A, Williams I, et al. CHoRUS Production Data Release V3. University of Virginia Dataverse; 2026. doi:10.18130/V3/ATG0DP ^53^
`o` Bensoussan Y, Sigaras A, Rameau A, et al. Bridge2AI-Voice: An ethically sourced, diverse voice dataset linked to health information. Published online May 1, 2026. doi:10.13026/8XBN-NQ66 ^54^
`o` AI-READI Consortium. Flagship Dataset of Type 2 Diabetes from the AI-READI Project. Published online 2025:3.87 TB, 358999 files. doi:10.60775/FAIRHUB.3 ^55^

### Evaluation Rubric

We provide an evaluation rubric and scoring methodology aligned with our AI-readiness criteria. A HTML scoring sheet was provided, and is noted below under Software.

`o` Clark T, Muñoz Torres M. Rubric for review of AI-readiness evaluation criteria. 2026 Sep 11. doi:10.5281/ZENODO.22702086 ^124^

### Software

`o` **FAIRSCAPE AI-readiness Framework**

`o` Niestroy J, Levinson M, Al Manir S, Clark T. FAIRSCAPE Documentation and Software. Zenodo 2026 Nov 8. https://doi.org/10.5281/zenodo.24062158. ^86^
`o` **RO-Crate Validation Classes**

`o` Niestroy, J., et al. 2026 - FAIRSCAPE Pydantic Models (v1.0.1). Zenodo. doi: 10.5281/zenodo.18234523 ^127^
`o` Leo, S., et al. 2025 - ro-crate-py (0.14.2). Zenodo. doi: 10.5281/zenodo.17342107 ^128^
`o` **FAIRSCAPE AI-readiness Scoring Sheet**

`o` Justin Niestroy. Fairscape AI-readiness Grader. Zenodo; 2026. doi:10.5281/ZENODO.23220189 ^85^

## Acknowledgements

This work was funded by National Institutes of Health Bridge2AI program awards OT2OD032742, OT2OD032644, OT2OD032720, OT2OD032701, U54HG012510, 5U54HG012517, and 5U54HG012513; and by the Frederick Thomas Fund of the University of Virginia. JNH has been supported with an EMBO Postdoctoral Fellowship (ALTF 556-2022).

We thank Carole Goble, Trey Ideker, Maryann Martone, and Randall Moorman for very helpful discussions, which improved this article.

## Use of Generative AI in the Writing Process

During revision, Google Gemini 3 Flash (December 2025) was used by the authors to flag passages where arguments were unclear or out of sequence, and to note any missing required sections. All rewriting and reorganization was performed by the authors. The manuscript includes no AI-generated text. The authors take full responsibility for the content.

The content of this article is solely the responsibility of the authors and does not necessarily represent the official views of the National Institutes of Health.

## Author Contributions

TC and MCMT conceived the study, oversaw the collaboration, standards alignment, and discussions, led development of the metadata architecture, and prepared the initial manuscript. TC and SAM, JN, and MAL developed the FAIRSCAPE RO-Crate packaging tools. TC and SAM developed the EVI profiles. JHC, MJ, JR, and MCMT developed the LinkML-based schema and LinkML AI-assist tools. JN, TC, and MCMT developed the criteria conformance rubric, scoring model, and scoring assist tools. TC, SAM, and JAP managed CM4AI metadata preparation and release testing. SSG, BP, AEW, and MAL managed metadata packaging for Bridge2AI clinical programs. TC prepared the final manuscript with review, discussion, and edits by MCMT, JAP, BP, SSG, and SJR. All authors contributed significantly to key discussions that informed the preparation, content, editing, and final review of the manuscript. All authors reviewed and approved the manuscript.

## REFERENCES

1. Topol EJ. High-performance medicine: the convergence of human and artificial intelligence. Nat Med. 2019;25(1):44–56. doi:10.1038/s41591-018-0300-7

2. Clissa L, Lassnig M, Rinaldi L. How big is Big Data? A comprehensive survey of data production, storage, and streaming in science and industry. Front Big Data. 2023;6:1271639. doi:10.3389/fdata.2023.1271639

3. Strickland E. Andrew Ng, AI Minimalist: The Machine-Learning Pioneer Says Small is the New Big. IEEE Spectr. 2022;59(4):22–50. doi:10.1109/MSPEC.2022.9754503

4. Walsh I, Fishman D, Garcia-Gasulla D, et al. DOME: recommendations for supervised machine learning validation in biology. Nat Methods. 2021;18(10):1122–1127. doi:10.1038/s41592-021-01205-4

5. Hiniduma K, Byna S, Bez JL. Data Readiness for AI: A 360-Degree Survey. ACM Comput Surv. 2025;57(9):1–39. doi:10.1145/3722214

6. Committee on Realizing Opportunities for Advanced and Automated Workflows in Scientific Research, Board on Research Data and Information, Board on Mathematical Sciences and Analytics, et al. Automated Research Workflows for Accelerated Discovery: Closing the Knowledge Discovery Loop. National Academies Press; 2022:26532. doi:10.17226/26532

7. Ng MY, Youssef A, Miner AS, et al. Perceptions of Data Set Experts on Important Characteristics of Health Data Sets Ready for Machine Learning: A Qualitative Study. JAMA Netw Open. 2023;6(12):e2345892. doi:10.1001/jamanetworkopen.2023.45892

8. Gitelman L, ed. “Raw Data” Is an Oxymoron. The MIT Press; 2013. doi:10.7551/mitpress/9302.001.0001

9. Leonelli S. Data Governance is Key to Interpretation: Reconceptualizing Data in Data Science. Harvard Data Science Review. Published online June 22, 2019. doi:10.1162/99608f92.17405bb6

10. Obermeyer Z, Powers B, Vogeli C, Mullainathan S. Dissecting racial bias in an algorithm used to manage the health of populations. Science. 2019;366(6464):447–453. doi:10.1126/science.aax2342

11. Sambasivan N, Kapania S, Highfill H, Akrong D, Paritosh P, Aroyo LM. “Everyone wants to do the model work, not the data work”: Data Cascades in High-Stakes AI. In: Proceedings of the 2021 CHI Conference on Human Factors in Computing Systems. ACM; 2021:1–15. doi:10.1145/3411764.3445518

12. Roberts M, Driggs D, Thorpe M, et al. Common pitfalls and recommendations for using machine learning to detect and prognosticate for COVID-19 using chest radiographs and CT scans. Nat Mach Intell. 2021;3(3):199–217. doi:10.1038/s42256-021-00307-0

13. Wilkinson MD, Dumontier M, Aalbersberg IjJ, et al. The FAIR Guiding Principles for scientific data management and stewardship. Scientific Data. 2016;3:160018. 10.1038/sdata.2016.18

14. National Institutes of Health. Bridge to Artificial Intelligence (Bridge2AI). Published online February 7, 2023. Accessed February 9, 2023. https://commonfund.nih.gov/bridge2ai

15. Chakir A, Andry JF, Ullah A, Bansal R, Ghazouani M, eds. Engineering Applications of Artificial Intelligence. Springer Nature Switzerland; 2024. doi:10.1007/978-3-031-50300-9

16. Russell SJ, Norvig P. Artificial Intelligence: A Modern Approach. Fourth edition. Pearson; 2021.

17. McCarthy J. A Proposal for the Dartmouth Summer Research Project on Artificial Intelligence. AI Magazine. 2006;27(4):12–14.

18. Gao S, Fang A, Huang Y, et al. Empowering biomedical discovery with AI agents. Cell. 2024;187(22):6125–6151. doi:10.1016/j.cell.2024.09.022

19. Kidwai-Khan F, Wang R, Skanderson M, Brandt CA, Fodeh S, Womack JA. A Roadmap to Artificial Intelligence (AI): Methods for Designing and Building AI ready Data for Women’s Health Studies. medRxiv. Published online May 30, 2023:2023.05.25.23290399. doi:10.1101/2023.05.25.23290399

20. Grobelnik M, Perset K, Russell S. What is AI? Can you make a clear distinction between AI and non-AI systems? Published online March 6, 2024. https://oecd.ai/en/wonk/definition

21. Regulation (EU) 2024/1689 of the European Parliament and of the Council of 13 June 2024 laying down harmonised rules on artificial intelligence (Artificial Intelligence Act). Official Journal of the European Union. 2024;(L1689). https://eur-lex.europa.eu/eli/reg/2024/1689/oj/eng

22. Altman RB, Levitt M. What is Biomedical Data Science and Do We Need an Annual Review of It? Annu Rev Biomed Data Sci. 2018;1(1):i-iii. doi:10.1146/annurev-bd-01-041718-100001

23. Shirey R. Internet Security Glossary, Version 2. Published online March 2, 2013. https://datatracker.ietf.org/doc/rfc4949/

24. Stevenson A, ed. Shorter Oxford English Dictionary on Historical Principles. 1: A - M / [Ed.: Angus Stevenson]. 6. ed. Oxford University Press; 2007.

25. NIST. NIST Computer Security Resource Center - Glossary. Published online June 6, 2024. https://csrc.nist.gov/glossary/term/data_element

26. W3C Schema.org Community Group. Schema.org: Dataset. Published online June 6, 2024.

27. Chaddad A, Peng J, Xu J, Bouridane A. Survey of Explainable AI Techniques in Healthcare. Sensors. 2023;23(2):634. doi:10.3390/s23020634

28. Juty N, Wimalaratne SM, Soiland-Reyes S, Kunze J, Goble CA, Clark T. Unique, Persistent, Resolvable: Identifiers as the foundation of FAIR. Data Intelligence. 2020;2(1-2):30–39. doi:10.5281/zenodo.3267434

29. Gil Y, Miles S, Belhajjame K, et al. PROV Model Primer: W3C Working Group Note 30 April 2013. Published online April 30, 2013. https://www.w3.org/TR/prov-primer/

30. ACD AI WG. Report of the Advisory Committee to the Director Working Group on AI. Published online December 6, 2019. https://www.acd.od.nih.gov/documents/presentations/12132019AI_FinalReport.pdf

31. U.S. Department of Health and Human Services. HIPAA Privacy Rule. 2026. https://www.ecfr.gov/current/title-45/subtitle-A/subchapter-C/part-164

32. U.S. Department of Health and Human Services. Federal Policy for the Protection of Human Subjects (Common Rule). https://www.ecfr.gov/current/title-45/subtitle-A/subchapter-A/part-46

33. European Parliament and Council of the European Union. General Data Protection Regulation. Official Journal of the European Union. 2016;L119:1–88

34. The National Commission for the Protection of Human Subjects of, Biomedical and Behavioral Research. The Belmont Report: Ethical Principles and Guidelines for the Protection of Human Subjects of Research. Published online April 18, 1979.

35. World Medical Association. World Medical Association Declaration of Helsinki: Ethical Principles for Medical Research Involving Human Participants. JAMA. 2025;333(1):71–74. doi:10.1001/jama.2024.21972

36. Akhtar M, Benjelloun O, Conforti C, et al. Croissant: A Metadata Format for ML-Ready Datasets. In: Proceedings of the Eighth Workshop on Data Management for End-to-End Machine Learning. 2024:1–6. doi:10.1145/3650203.3663326

37. Jain N, Akhtar M, Giner-Miguelez J, et al. A Standardized Machine-readable Dataset Documentation Format for Responsible AI. arXiv. Preprint posted online June 4, 2024:arXiv:2407.16883. doi:10.48550/arXiv.2407.16883

38. Masic I, Muhamedagic B, Kulasin I, Masic Z, Sanousi A. Health Level Seven (hl 7): Short Overview. Mater Sociomed. 2009;21(3):170. doi:10.5455/msm.2009.21.170-174

39. Gebru T, Morgenstern J, Vecchione B, et al. Datasheets for datasets. Commun ACM. 2021;64(12):86–92. doi:10.1145/3458723

40. Chmielinski KS, Newman S, Taylor M, et al. The Dataset Nutrition Label (2nd Gen): Leveraging Context to Mitigate Harms in Artificial Intelligence. arXiv. Preprint posted online March 10, 2022:arXiv:2201.03954. Accessed November 2, 2023. http://arxiv.org/abs/2201.03954

41. Majithia N, Carey-Wilson T, Simperl E. A Framework for AI-Ready Data. 2025. doi:10.61557/ODEX6433

42. Majithia N, Carey-Wilson T, Simperl E, Shadbolt N. An actionable framework for AI-ready data. AI Magazine. 2026;47(1):e70054. doi:10.1002/aaai.70054

43. Clark T, Caufield H, Parker JA, et al. AI-readiness for Biomedical Data: Bridge2AI Recommendations. bioRxiv. Published online November 24, 2024:2024.10.23.619844. doi:10.1101/2024.10.23.619844

44. Al Manir S, Clark T. Crosswalk of AI-readiness criteria: Clark et al. 2026 and Majithia et al. 2026. Published online October 12, 2026. doi:10.5281/ZENODO.23306249

45. Paullada A, Raji ID, Bender EM, Denton E, Hanna A. Data and its (dis)contents: A survey of dataset development and use in machine learning research. Patterns. 2021;2(11):100336. doi:10.1016/j.patter.2021.100336

46. Leonelli S, Tempini N, eds. Data Journeys in the Sciences. Springer International Publishing; 2020. doi:10.1007/978-3-030-37177-7

47. Leonelli S. Data-Centric Biology: A Philosophical Study. The University of Chicago Press; 2016.

48. Edwards PN. A Vast Machine: Computer Models, Climate Data, and the Politics of Global Warming. MIT Press; 2010.

49. Bowker GC, Star SL. Sorting Things out: Classification and Its Consequences. MIT Press; 2000.

50. Latour B. Pandora’s Hope: Essays on the Reality of Science Studies. 2. print. Harvard University Press; 2000.

51. Latour B, Woolgar S. Laboratory Life: The Construction of Scientific Facts. Princeton University Press; 1986.

52. Clark T, Parker J, Al Manir S, et al. Cell Maps for Artificial Intelligence - June 2026 Data Release (Beta). Published online July 15, 2026:4067695317, 4497827175, 4974378970, 139066, 175962, 75070, 30921, 1203765, 96135, 115990, 4063091401, 4966882033, 4492541281. doi:10.18130/V3/HIGT4C

53. Rosenthal E, Kamaleswaran R, Levites Strekalova YA, et al. CHoRUS Production Data Release V3. Published online October 29, 2026:102156. doi:10.18130/V3/ATG0DP

54. Bensoussan Y, Sigaras A, Rameau A, et al. Bridge2AI-Voice: An ethically-sourced, diverse voice dataset linked to health information. Published online May 1, 2026. doi:10.13026/8XBN-NQ66

55. AI-READI Consortium. Flagship Dataset of Type 2 Diabetes from the AI-READI Project. Published online 2025:3.87 TB, 358999 files. doi:10.60775/FAIRHUB.3

56. Artificial intelligence — Data quality for analytics and machine learning (ML). Published online 2024:ISO/iEC 5259. https://www.iso.org/standard/81088.html

57. International Standards Organiztion. ISO/IEC 42001:2023(E). Published online 2023.

58. Lebo T, Sahoo S, McGuinness D, et al. PROV-O: The PROV Ontology W3C Recommendation 30 April 2013. Published online 2013. http://www.w3.org/TR/prov-o/

59. Al Manir S, Niestroy J, Levinson MA, Clark T. Evidence Graphs: Supporting Transparent and FAIR Computation, with Defeasible Reasoning on Data, Methods, and Results. In: Glavic B, Braganholo V, Koop D, eds. Provenance and Annotation of Data and Processes. Vol 12839. Lecture Notes in Computer Science. Springer International Publishing; 2021:39–50. doi:10.1007/978-3-030-80960-7_3

60. Al Manir S, Niestroy J, Levinson M, Clark T. EVI: The Evidence Graph Ontology, OWL 2 Vocabulary. Published online March 16, 2026. 10.5281/zenodo.4630931

61. ISO/TC 276. Biotechnology — Provenance information model for biological material and data — Part 2: Common provenance model. Published online June 2026:ISO 23494:2(2026). https://www.iso.org/standard/87714.html

62. Moreau L, Missier P, Belhajjame K, et al. PROV-DM: The PROV Data Model: W3C Recommendation 30 April 2013. World Wide Web Consortium; 2013. http://www.w3.org/TR/prov-dm/

63. Software engineering — Software product Quality Requirements and Evaluation (SQuaRE) — Data quality model. Published online 2008:ISO/IEC 25012. https://www.iso.org/standard/35736.html

64. Systems and software engineering — Systems and software Quality Requirements and Evaluation (SQuaRE) — Measurement of data quality. Published online 2015:ISO/IEC 25024. https://www.iso.org/standard/35749.html

65. Information technology — Artificial intelligence — Guidance on risk management. Published online 2023:ISO/IEC 23894:2023. https://www.iso.org/standard/77304.html

66. Regulation (EU) 2026/1744 of the European Parliament and of the Council of 8 July 2026 amending Regulations (EU) 2024/1689, (EU) 2018/1139 and (EU) 2023/1230 as regards the simplification of the implementation of harmonised rules on artificial intelligence (Digital Omnibus on AI). Official Journal of the European Union. 2026;((L1744)). https://eur-lex.europa.eu/legal-content/EN/TXT/?uri=OJ:L_202601744

67. Shapin S. Pump and Circumstance: Robert Boyle’s Literary Technology. In: Hellyer M, ed. The Scientific Revolution. Blackwell; 2003.

68. Callaghan S, Donegan S, Pepler S, et al. Making Data a First Class Scientific Output: Data Citation and Publication by NERC’s Environmental Data Centres. IJDC. 2012;7(1):107–113. doi:10.2218/ijdc.v7i1.218

69. National Academies of Science, Engineering, and Medicine. Reproducibility and Replicability in Science. THE NATIONAL ACADEMIES PRESS; 2019. doi:10.17226/25303

70. Ciccarese P, Soiland-Reyes S, Belhajme K, Gray AJ, Goble C. PAV ontology: Provenance, Authoring and Versioning. Journal of Biomedical Semantics. 2013;4(37). doi:10.1186/2041-1480-4-37

71. Begley CG, Ioannidis JPA. Reproducibility in Science: Improving the Standard for Basic and Preclinical Research. Circulation Research. 2015;116(1):116–126. doi:10.1161/CIRCRESAHA.114.303819

72. Clark T, Mohan J, Schaffer L, et al. Cell Maps for Artificial Intelligence: AI-Ready Maps of Human Cell Architecture from Disease-Relevant Cell Lines. Preprint posted online May 24, 2024. doi:10.1101/2024.05.21.589311

73. Lenkiewicz J, Churas C, Hu M, et al. Cell Mapping Toolkit: an end-to-end pipeline for mapping subcellular organization. Bioinformatics. 2025;41(6):btaf205. doi:10.1093/bioinformatics/btaf205

74. Schaffer LV, Hu M, Qian G, et al. Multimodal cell maps as a foundation for structural and functional genomics. Nature. Published online April 9, 2025. doi:10.1038/s41586-025-08878-3

75. Mitchell M, Wu S, Zaldivar A, et al. Model Cards for Model Reporting. Proceedings of the Conference on Fairness, Accountability, and Transparency. Published online January 29, 2019:220–229. doi:10.1145/3287560.3287596

76. Crisan A, Drouhard M, Vig J, Rajani N. Interactive Model Cards: A Human-Centered Approach to Model Documentation. In: 2022 ACM Conference on Fairness, Accountability, and Transparency. 2022:427–439. doi:10.1145/3531146.3533108

77. Adashi EY, Walters LB, Menikoff JA. The Belmont Report at 40: Reckoning With Time. Am J Public Health. 2018;108(10):1345–1348. doi:10.2105/AJPH.2018.304580

78. Bailey M, Dittrich D, Kenneally E, Maughan D. The Menlo Report. IEEE Secur Privacy Mag. 2012;10(2):71–75. doi:10.1109/MSP.2012.52

79. Carroll SR, Garba I, Figueroa-Rodríguez OL, et al. The CARE Principles for Indigenous Data Governance. Data Science Journal. 2020;19:43. doi:10.5334/dsj-2020-043

80. Carroll MW. Creative Commons and the New Intermediaries. Mich St L Rev. 2006;45. http://works.bepress.com/michael_carroll/1

81. Lin D, Crabtree J, Dillo I, et al. The TRUST Principles for digital repositories. Sci Data. 2020;7(1):144. doi:10.1038/s41597-020-0486-7

82. Justin Niestroy, Al Manir MS, Levinson M, Munoz-Torres M, Clark T. Fairscape RO-Crate Profile. Published online October 8, 2026. doi:10.5281/ZENODO.23243591

83. Joacimiak M, Caufield H, Banks O, et al. Bridge2AI Datasheets for Datasets: Schema. Published online December 27, 2025. https://github.com/bridge2ai/data-sheets-schema

84. Moxon SAT, Solbrig H, Harris NL, et al. LinkML: An Open Data Modeling Framework. GigaScience. Published online December 12, 2025:giaf152. doi:10.1093/gigascience/giaf152

85. Justin Niestroy, Clark T, Munoz-Torres M. AI-Readiness Grader. Published online October 7, 2026. doi:10.5281/ZENODO.23220189

86. Justin Niestroy, Levinson M, Al Manir MS, Clark T. fairscape/fairscape.github.io: FAIRSCAPE Documentation. Published online November 8, 2026. doi:10.5281/ZENODO.24062158

87. Al Manir S, Levinson MA, Niestroy J, et al. The FAIRSCAPE AI-readiness Framework for Biomedical Research. BioRxiv. Preprint posted online March 4, 2026. doi:10.1101/2024.12.23.629818

88. Thomas DM, Knight R, Gilbert JA, et al. Transforming Big Data into AI-ready data for nutrition and obesity research. Obesity. 2024;32(5):857–870. doi:10.1002/oby.23989

89. Poduval B, McPherron RL, Walker R, et al. AI-ready data in space science and solar physics: problems, mitigation and action plan. Front Astron Space Sci. 2023;10:1203598. doi:10.3389/fspas.2023.1203598

90. Lindemann T, Häberlein L. Contours of a research ethics and integrity perspective on open science. Front Res Metr Anal. 2023;8:1052353. doi:10.3389/frma.2023.1052353

91. Committee on Responsible Science, Committee on Science, Engineering, Medicine, and Public Policy, Policy and Global Affairs, National Academies of Sciences, Engineering, and Medicine. Fostering Integrity in Research. National Academies Press; 2017:21896. doi:10.17226/21896

92. The FAIRsharing Team. FAIRsharing.org. Published online 2024. Accessed October 30, 2024. https://fairsharing.org/

93. National Institutes of Health O of the D. Generalist Repository Ecosystem Initiative. Published online July 6, 2023. https://datascience.nih.gov/data-ecosystem/generalist-repository-ecosystem-initiative

94. Riccardo Albertoni, David Browning, Simon JS Cox, Alejandra Gonzalez Beltran, Andrea Perego, Peter Winstanley. Data Catalog Vocabulary (DCAT) - Version 3. Published online August 22, 2024. https://www.w3.org/TR/vocab-dcat-3/

95. Guha RV, Brickley D, Macbeth S. Schema.org: evolution of structured data on the web. Communications of the ACM. 2016;59(2):44–51. doi:10.1145/2844544

96. DataCite Metadata Working Group. DataCite Metadata Schema Documentation for the Publication and Citation of Research Data and Other Research Outputs v4.7. Ashton J, Bernal I, Devaraju A, et al., eds. Published online March 3, 2026. doi:10.14454/QDD3-PS68

97. Brickley D, Guha RV. RDF Vocabulary Description Language 1.0: RDF Schema. World Wide Web Consortium; 2004. http://www.w3.org/TR/rdf-schema/

98. Sporny M, Longley D, Kellogg G, Lanthaler M, Champin PA, Lindström N. JSON-LD 1.1: A JSON-based Serialization for Linked Data. Kellogg G, Champin PA, Longley D, eds. W3C Recommendation 16 July 2020. Published online July 16, 2020. https://www.w3.org/TR/json-ld/

99. Le Dévédec SE, Hettne K, Gambardella A, Schultes E, Posth S. FAIA - FAIR AI Attribution White Paper. Published online June 17, 2026. doi:10.5281/ZENODO.20083769

100. Observational Medical Outcomes Partnership (OMOP). Standardized Data: The OMOP Common Data Model. Published online 2024. https://www.ohdsi.org/data-standardization/

101. Bandrowski AE, Martone ME. RRIDs: A Simple Step toward Improving Reproducibility through Rigor and Transparency of Experimental Methods. Neuron. 2016;90(3):434–436. doi:10.1016/j.neuron.2016.04.030

102. European Organization For Nuclear Research, OpenAIRE. Zenodo. Published online 2013. doi:10.25495/7GXK-RD71

103. Troupin C, Muñoz C, Fernández JG, Rújula MA. Scientific results traceability: software citation using GitHub & Zenodo. In: Proceedings of the International Conference on Marine Data and Information Systems. 2018:4.

104. Potter M, Smith T. Making code citable with Zenodo and GitHub. Published online July 25, 2015. https://www.software.ac.uk/blog/making-code-citable-zenodo-and-github

105. Abramatic JF, Di Cosmo R, Zacchiroli S. Building the universal archive of source code. Commun ACM. 2018;61(10):29–31. doi:10.1145/3183558

106. Blischak JD, Davenport ER, Wilson G. A Quick Introduction to Version Control with Git and GitHub. Ouellette F, ed. PLoS Comput Biol. 2016;12(1):e1004668. doi:10.1371/journal.pcbi.1004668

107. Akers KG, Sarkozy A, Wu W, Slyman A. ORCID Author Identifiers: A Primer for Librarians. Medical Reference Services Quarterly. 2016;35(2):135–144. doi:10.1080/02763869.2016.1152139

108. Lammey R. Solutions for identification problems: a look at the Research Organization Registry. Sci Ed. 2020;7(1):65–69. doi:10.6087/kcse.192

109. The pandas development team. pandas-dev/pandas: Pandas. Published online June 28, 2026. doi:10.5281/ZENODO.3509134

110. Ritchie Vink, Alexander Beedie, nameexhaustion, et al. pola-rs/polars: Python Polars 1.42.1. Published online June 30, 2026. doi:10.5281/ZENODO.21056103

111. Fowler D, Barratt J, Walsh P. Frictionless Data: Making Research Data Quality Visible. IJDC. 2018;12(2):274–285. doi:10.2218/ijdc.v12i2.577

112. Wright, A., Andrews, A., Hutton, B., Dennis, G. JSON Schema: A Media Type for Describing JSON Documents. Published online June 16, 2022. https://json-schema.org/draft/2020-12/json-schema-core

113. Pydantic. Pydantic: Documentation for version: v2.13.4. https://docs.pydantic.dev/latest/

114. Albertoni R, Isaac A. W3C Data Quality Vocabulary (DQV). Published online December 15, 2016.

115. Rostamzadeh N, Mincu D, Roy S, et al. Healthsheet: Development of a Transparency Artifact for Health Datasets. In: 2022 ACM Conference on Fairness, Accountability, and Transparency. ACM; 2022:1943–1961. doi:10.1145/3531146.3533239

116. Lawson J, Cabili MN, Kerry G, et al. The Data Use Ontology to streamline responsible access to human biomedical datasets. Cell Genomics. 2021;1(2):100028. doi:10.1016/j.xgen.2021.100028

117. National Institute of Standards and Technology (US). Secure Hash Standard. National Institute of Standards and Technology (U.S.); 2015:NIST FIPS 180–4. doi:10.6028/NIST.FIPS.180-4

118. Merkle RC. A Digital Signature Based on a Conventional Encryption Function. In: Pomerance C, ed. Advances in Cryptology — CRYPTO ‘87. Vol 293. Lecture Notes in Computer Science. Springer Berlin Heidelberg; 1988:369–378. doi:10.1007/3-540-48184-2_32

119. Health Level 7. FHIR Specification v5.0.0: R5. Published online March 26, 2023. https://hl7.org/fhir/index.html

120. Global Alliance for Genomics and Health (GA4GH). Data Access Committee Guiding Principles and Procedural Standards Policy. Published online 2021.

121. HL7. HL7 Terminology 6.1.0 - Confidentiality. Published online November 17, 2024. https://terminology.hl7.org/CodeSystem-v3-Confidentiality.html

122. RO-Crate Community. Research Object Crate (RO-Crate). Published online October 26, 2023. https://www.researchobject.org/ro-crate/

123. Jones MB, Boettiger C, Mayes AC, et al. CodeMeta: an exchange schema for software metadata. Published online 2023. https://w3id.org/codemeta/3.0

124. Clark T, Muñoz Torres M. Rubric for review of AI-readiness evaluation criteria. Published online September 11, 2026. doi:10.5281/ZENODO.22702086

125. Sefton P, Ó Carragáin E, Soiland-Reyes S, et al. RO-Crate Metadata Specification 1.2.0. Sefton P, Soiland-Reyes S, eds. researchobject.org. Preprint posted online June 4, 2025. doi:10.5281/ZENODO.13751027

126. World Health Organization. Ethics and Governance of Artificial Intelligence for Health: WHO Guidance. World Health Organization; 2021:150. https://www.who.int/publications/i/item/9789240029200

127. Justin Niestroy, Max Levinson, Sadnan Al Manir, Clark T. Fairscape Pydantic Models. Published online January 13, 2026. doi:10.5281/ZENODO.18234523

128. Simone Leo, Stian Soiland-Reyes, Ignacio Eguinoa, et al. ro-crate-py. Published online October 13, 2025. doi:10.5281/ZENODO.17342107

